# A zinc finger transcription factor tunes social behaviors by controlling transposable elements and immune response in prefrontal cortex

**DOI:** 10.1101/2023.04.03.535374

**Authors:** Natalie L. Truby, R. Kijoon Kim, Gabriella M. Silva, Xufeng Qu, Joseph A. Picone, Rebecca Alemu, Rachael L. Neve, Xiaohong Cui, Jinze Liu, Peter J. Hamilton

**Author notes:** These authors contributed equally to this work (Natalie L. Truby, R. Kijoon Kim, Gabriella M. Silva).

## Abstract

The neurobiological origins of social behaviors are incompletely understood. Here we utilized synthetic biology approaches to reprogram the function of ZFP189, a transcription factor whose expression and function in the rodent prefrontal cortex was previously determined to be protective against stress-induced social deficits. We created novel synthetic ZFP189 transcription factors including ZFP189^VPR^, which activates the transcription of target genes and therefore exerts opposite functional control from the endogenous, transcriptionally repressive ZFP189^WT^. Upon viral delivery of these synthetic ZFP189 transcription factors to mouse prefrontal cortex, we observe that ZFP189-mediated transcriptional control promotes mature dendritic spine morphology on transduced pyramidal neurons. Interestingly, dysregulation of ZFP189-mediated transcription in this brain area, achieved by delivery of synthetic ZFP189^VPR^, precipitates social behavioral deficits in terms of social interaction, motivation, and the cognition necessary for the maintenance of social hierarchy, without other observable behavioral deficits. By performing RNA sequencing in virally manipulated prefrontal cortex tissues, we discover that ZFP189 transcription factors of opposing regulatory function have opposite influence on the expression of genetic transposable elements as well as genes that participate in immune functions. Collectively, this work reveals that ZFP189 function in the prefrontal cortex coordinates structural and transcriptional neuroadaptations necessary for social behaviors by binding transposable element-rich regions of DNA to regulate immune-related genes. Given the evidence for a co-evolution of social behavior and the brain immune response, we posit that ZFP189 may have evolved to augment brain transposon-associated immune function as a way of enhancing an animal’s capacity for functioning in social groups.

## Introduction

Krüppel-associated box (KRAB) zinc finger proteins (KZFPs) represent an evolutionarily ancient class of protein (1,2) and the largest family of transcription factors (TFs) encoded in the mammalian genome (3). While growing evidence suggests that many KZFPs play a role in regulating genomic transposable elements (TEs) as well as controlling the expression of protein coding genes (4,5), the complete gene-regulatory functions of individual KZFPs and how their function within certain brain areas contribute to neuroadaptations and organismal behavior remains poorly understood.

Earlier work employing co-expression analyses of RNA-sequencing (RNAseq) datasets of limbic brain areas from mice subjected to chronic social stress identified one member of the KZFP gene family, *Zfp189*, as the top regulatory transcript responsible for manifesting transcriptional networks in the prefrontal cortex (PFC) unique to stress ‘resilient’ rodents (6). These resilient animals were classified as possessing the capacity to endure chronic social stress without developing detectable behavioral deficits in social interaction and other stress-related phenotypes. Viral over-expression or CRISPR-mediated activation of *Zfp189* mRNA expression in PFC rescued social deficits in stress-exposed mice (6), establishing PFC *Zfp189* as a molecular mediator of resistance to stress-induced social deficits.

Indeed, social dysfunction manifests in a multitude of neurological disorders including schizophrenia, autism spectrum disorder, and Williams syndrome (7) and in response to stressful life events which contribute risk for developing neuropsychiatric syndromes like major depressive disorder and post-traumatic stress disorder (8). However, the brain molecular mechanisms that govern social behavior, and how these mechanisms are impacted in disease states, are not fully understood.

Here, to interrogate the gene targets and nuanced behaviors controlled by ZFP189, we developed novel synthetic ZFP189 TFs, each capable of exerting distinct forms of transcriptional control at *in vivo* ZFP189 target genes. We replaced the endogenous repressive KRAB moiety of ZFP189 with the transcriptional activator VP64-p65-Rta (VPR) to create the novel synthetic TF ZFP189^VPR^. We combined PFC viral delivery of these synthetic TFs with transcriptome profiling and behavioral assays to uncover the molecular mechanisms by which ZFP189 governs specific social behaviors. Virally delivering these ZFP189 TFs to the mouse PFC resulted in maturation of mature dendritic spines on pyramidal neurons. We were surprised to discover that synthetically inverting the transcriptional control exerted by ZFP189 within PFC precipitated pronounced social deficits via increased transcription of TEs and decreased expression of interferon pathogen responsive genes. Among other social deficits, both socially dominant and subordinate ZFP189^VPR^-treated mice prove unable to maintain their established position in a social hierarchy, suggesting that ZFP189-regulated neurobiological mechanisms normally function to enable the social cognition necessary for participation in a social group. This body of work collectively indicates that ZFP189 tunes an animal’s proclivity for social behavior by orchestrating a TE-regulated adaptive immune response in PFC cortical neurons to drive neuroplasticity and complex social behaviors.

## Results

### Synthetic ZFP189 transcription factors distinctly regulate gene expression and neuronal morphology

To investigate the direct gene-regulatory functions of ZFP189, we employed a synthetic biology approach wherein we created three synthetic ZFP189 TFs: ZFP189^WT^ which is identical to the endogenously expressed ZFP189 protein and contains the wild-type N-terminal KRAB domain and a C-terminal Cys_2_-His_2_ DNA-binding domain; ZFP189^VPR^ wherein the endogenous KRAB domain is replaced with the synthetic transcriptional activator VPR; and ZFP189^DN^ in which any transcriptional regulatory domain is removed, which serves to control for any non-specific effects of the expression vector or the over-expressed ZFP189 DNA-binding domain (Fig. 1A). Since the DNA binding motifs of mouse ZFP189 have not been determined, we inserted the experimentally determined (9) DNA response element (RE) motifs of the human ortholog ZNF189, which is 92% identical to ZFP189 on the amino acid level, upstream of the thymidine kinase (TK) promoter to drive luciferase expression as a gene reporter for ZFP189 function (Fig. 1B). By co-transfecting mouse neuroblastoma Neuro-2a (N2a) cells with both the ZFP189 RE luciferase plasmid and iterations of our ZFP189 TFs, we are able to observe that ZFP189^VPR^ induces robust targeted gene activation, ZFP189^WT^ induces gene repression, and ZFP189^DN^ exerts no regulatory control (Fig. 1C). The null effect of ZFP189^DN^ supports its use as the most appropriate control in future studies. By co-transfecting ZFP189^VPR^ alongside the other ZFP189 TF variants, we observe competition for regulation of luciferase expression. ZFP189^DN^ impedes the gene activation of ZFP189^VPR^ likely via steric interference and competition for ZFP189 REs at the luciferase promoter, whereas ZFP189^WT^ further decreases ZFP189^VPR^ function, presumably via the additional action of the repressive KRAB domain (Fig. 1D). Lastly, to investigate the requirement of DNA-binding to putative ZFP189 REs in these TF functions, we excised the promoter ZFP189 RE motifs in our luciferase plasmid (ZFP189 null) and compared the gene-regulatory functions of each ZFP189 TF. In the absence of ZFP189 DNA binding sites, we observe a 93% reduction in the gene-activating function of ZFP189^VPR^ (Fig. 1E). Conversely, we observe an increase in luciferase expression with ZFP189^WT^, indicating the loss of ZFP189 binding releases the repressive ZFP189^WT^ function (Fig. 1F). Interestingly, a similar effect is observed in N2a cells when no synthetic ZFP189 TF is co-transfected (Supp. Fig. 1), indicating the endogenous presence of ZFP189-like protein that functions similarly to our ZFP189^WT^. Lastly, the availability of ZFP189 DNA binding sites did not alter ZFP189^DN^ function, further indicating its lack of transcriptional control (Fig. 1G). Collectively, these data illustrate that our synthetic ZFP189 TFs can either up-or down-regulate the expression of target genes and this is dependent on direct binding to genomic ZFP189 RE DNA motifs.

**Figure 1:**
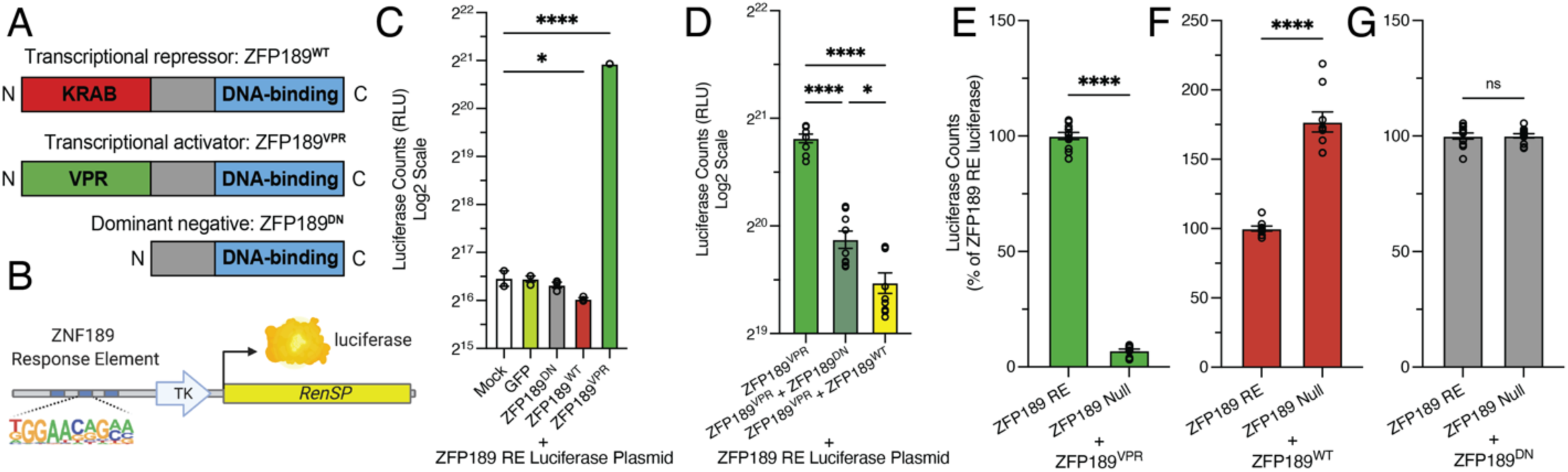
Synthetic ZFP189 transcription factors exert opposite transcriptional control at a luciferase target gene *in vitro*. **(A)** A cartoon representation of the three synthetic ZFP189 transcription factor (TF) proteins. **(B)** A cartoon of the ZFP189 DNA response element (RE) luciferase plasmid reporter target gene. TK is thymidine kinase gene promoter. *RenSP* is the LightSwitch™ luciferase gene. **(C)** Transient transfection of the ZFP189 RE luciferase plasmid and individual ZFP189 TFs in N2a cells reveals opposing transcriptional regulation of the luciferase target gene. Specifically, ZFP189^WT^ down-regulates and ZFP189^VPR^ up-regulates luciferase-driven relative light units (RLUs) relative to control conditions. One-way ANOVA followed by Bonferroni’s multiple comparisons test relative to mock transfection condition. * p-value < 0.05, **** p-value < 0.0001; n = 3 per condition. **(D)** Co-transfecting synthetic ZFP189 TFs of opposing function compete for regulation of the luciferase target gene. ZFP189^DN^, and to a greater degree ZFP189^WT^, inhibit ZFP189^VPR^-mediated gene activation. One-way ANOVA followed by Bonferroni’s multiple comparisons test. * p-value < 0.05, **** p-value < 0.0001; n = 9 per condition. **(E)** Removing ZFP189 REs from the promoter of our luciferase target gene (ZFP189 null) ablates the gene activation of ZFP189^VPR^. Data normalized to ZFP189 RE condition, by experiment. Two-tailed, unpaired Student’s *t*-test; **** p-value < 0.0001. n = 12 per condition. **(F)** Removing ZFP189 REs from the promoter of our luciferase target gene releases the gene repression of ZFP189^WT^. Data normalized to ZFP189 RE condition, by experiment. Two-tailed, unpaired Student’s *t*-test; **** p-value < 0.0001. n = 9 per condition. **(G)** Removing ZFP189 REs from the promoter of our luciferase target gene does not impact the functions of ZFP189^DN^. Data normalized to ZFP189 RE condition, by experiment. Two-tailed, unpaired Student’s *t*-test; p-value > 0.05. n = 12 per condition.

In the course of performing this research, it became apparent that N2a cells expressing the synthetic ZFP189^VPR^ exhibited more complex cellular morphologies, including an increased number and length of cellular outgrowths (Supp. Fig. 2). This hints that the unique transcriptional control exerted by ZFP189^VPR^ regulates the cellular programs governing cell growth and differentiation in the mouse neuronal lineage N2a cells.

**Figure 2:**
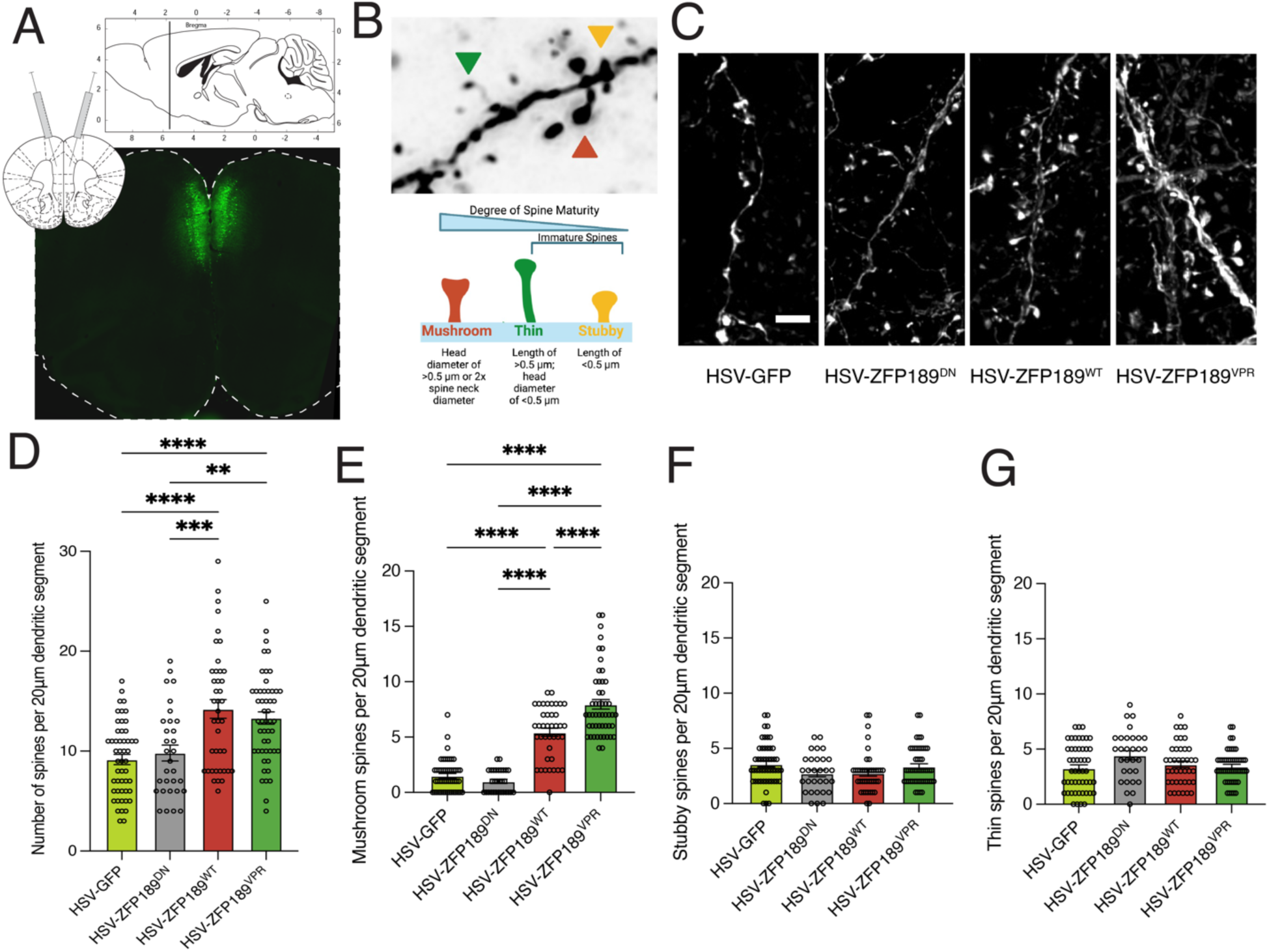
Viral delivery of synthetic ZFP189 transcription factors promote mature spine morphology in pyramidal neurons of the prefrontal cortex. **(A)** Representative image of viral targeting to the mouse prefrontal cortex (PFC). **(B)** Representative image of the three quantified spine morphologies (top) and the criteria for categorizing spines (bottom). **(C)** Representative pyramidal neuron dendritic segments from PFC tissues expressing GFP or one of the synthetic ZFP189 TFs. Scale bar is 2 μm. **(D)** Both HSV-ZFP189^WT^ and HSV-ZFP189^VPR^ (i.e., the transcriptionally functional ZFP189 TFs) similarly drive an increase in the number of spines per 20 μm dendritic segment. One-way ANOVA followed by Bonferroni’s multiple comparisons test. ** p-value < 0.01, *** p-value < 0.001, **** p-value < 0.0001; n = 10 segments from 3-4 mice per condition. **(E)** HSV-ZFP189^WT^ and, to a greater degree, HSV-ZFP189^VPR^ increase the number of mature mushroom spines on transduced pyramidal neurons of the PFC. One-way ANOVA followed by Bonferroni’s multiple comparisons test. **** p-value < 0.0001; n = 10 segments from 3-4 mice per condition. **(F)** No treatment affects the number of immature stubby spines on transduced neurons. One-way ANOVA followed by Bonferroni’s multiple comparisons test. p-value > 0.05; n = 10 segments from 3-4 mice per condition. **(G)** No treatment affects the number of immature thin spines on transduced neurons. One-way ANOVA followed by Bonferroni’s multiple comparisons test. p-value > 0.05; n = 10 segments from 3-4 mice per condition.

To explore this possibility further in the PFC, the brain area in which we identified *Zfp189* (6), we packaged each of our synthetic ZFP189 TFs into herpes simplex virus (HSV) vectors for stereotaxic viral delivery to awake and behaving mice. Upon delivery to the PFC (Fig. 2A-C), we observed that, despite exerting opposing forms of transcriptional control at *in vitro* target genes (Fig. 1C), both ZFP189^WT^ and ZFP189^VPR^ similarly increase the density of dendritic spines on transduced pyramidal neurons. Specifically, ZFP189^WT^, and to a greater degree, ZFP189^VPR^ increase the density of mature mushroom spines, with no effect on the immature stubby or thin spines (Fig. 2E-G). These results indicate that either transcriptionally functional ZFP189 TF regulates the neuroplasticity mechanisms that drive dendritic spine maturity on PFC pyramidal neurons.

### Inverting ZFP189-mediated transcriptional control in prefrontal cortex induces social behavioral deficits

We next sought to uncover the behavioral consequences of dysregulating ZFP189-mediated transcriptional control in the PFC. As PFC *Zfp189* expression was discovered to induce resilience to social stress-induced behavioral deficits, (6) we first virally delivered the synthetic ZFP189 TFs to the PFC, subjected the animals to a micro-defeat sub-threshold social stress, and performed social interaction and elevated plus maze behavioral testing (Fig. 3A), as we have previously(6,10,11). As expected, this mild stressor had no effect on the social behaviors of HSV-ZFP189^DN^ or HSV-ZFP189^WT^ treated mice (Fig. 3B). Much to our surprise, mice that received HSV-ZFP189^VPR^ in PFC showed social avoidance following the mild stressor (Fig. 3B). No viral treatment affected basal locomotion (Fig. 3C) or exploration of the open arm of an elevated plus maze (Fig. 3D). This indicates that inversion of the endogenous ZFP189 transcriptional control in PFC with ZFP189^VPR^ ablates social approach behaviors, without affecting basal or other anxiety-related behaviors.

**Figure 3:**
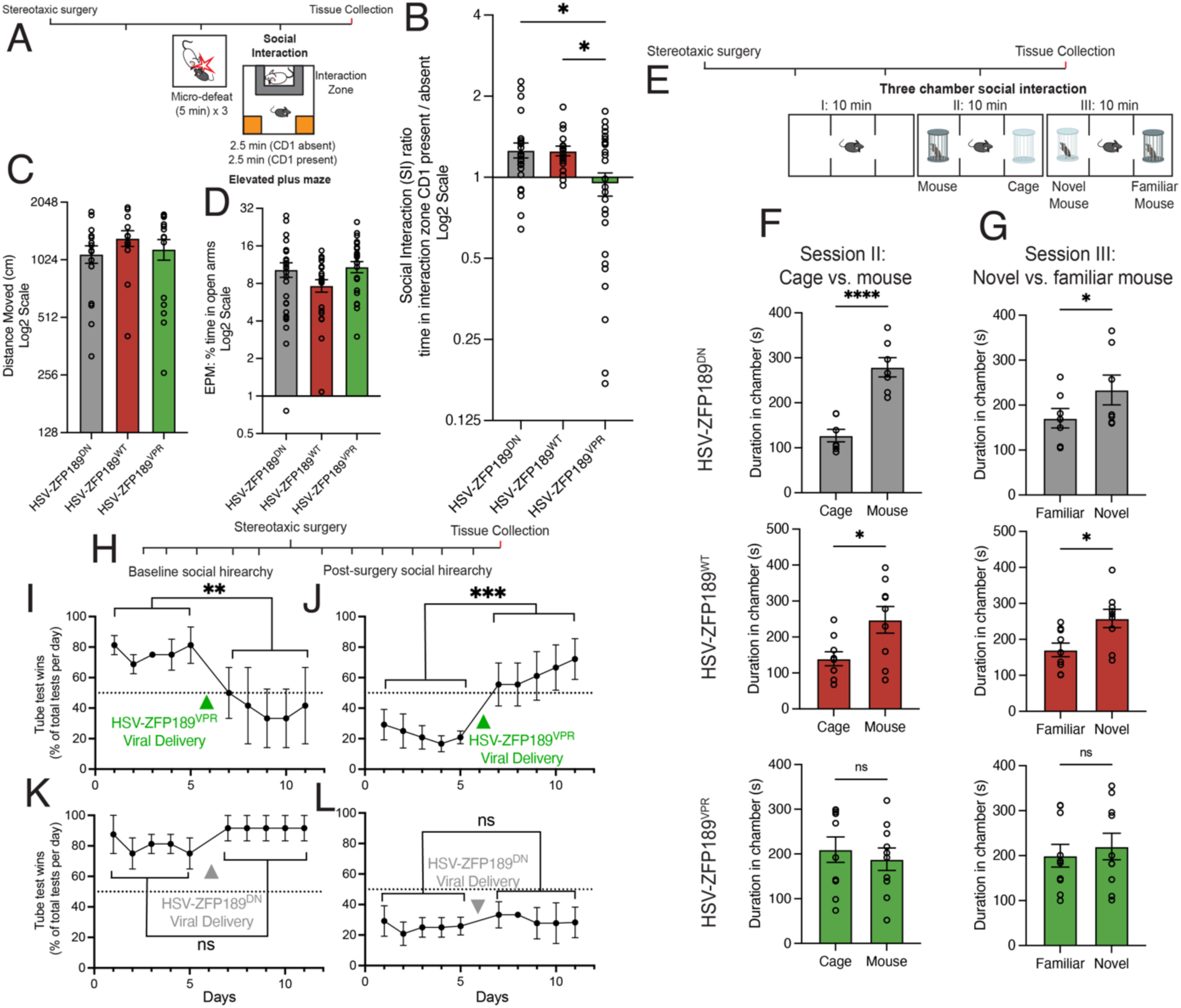
Inverting the natural transcriptional control of ZFP189 in the PFC with HSV-ZFP189^VPR^ ablates social function in mice. **(A)** Experimental timeline for viral delivery, micro-defeat, social interaction, and elevated plus maze testing. **(B)** Mice treated with HSV-ZFP189^VPR^ spend less time interacting with a CD1 social target mouse. Social interaction (SI) ratio quantifying the time test mice spent in interaction zone when social target mouse cage is empty relative to when the cage is occupied with novel CD1 mouse. Mice treated with HSV-ZFP189^DN^ or HSV-ZFP189^WT^ prefer to socialize (SI ratio >1), whereas HSV-ZFP189^VPR^ mice do not (SI ratio <1). One-way ANOVA followed by Bonferroni’s multiple comparisons test. * p-value < 0.05; n = 18 mice (HSV-ZFP189^DN^), n = 25 mice (HSV-ZFP189^WT^), n = 28 mice (HSV-ZFP189^VPR^). **(C)** No viral treatment condition affected locomotor behaviors in the SI test, measured in the first 2.5 min session when the target CD1 was absent. **(D)** No viral treatment condition affected time spent on the open arm of the elevated plus maze. One-way ANOVA followed by Bonferroni’s multiple comparisons test. p-value > 0.05. **(E)** Experimental timeline for three-chamber social interaction test. **(F)** In session II, when given the choice between interacting with an empty cage or a novel mouse, HSV-ZFP189^DN^ and HSV-ZFP189^WT^ treated mice spend more time interacting with the novel mouse, whereas HSV-ZFP189^VPR^ treated mice do not. Unpaired two-tailed Student’s *t*-test. **** p-value < 0.0001, * p-value < 0.05, ns p-value > 0.05. **(G)** Subjecting these same animals to session III wherein test mice are given a choice between interacting with the caged mouse from the previous round (Familiar) or a novel mouse, HSV-ZFP189^DN^ and HSV-ZFP189^WT^ treated mice again spend more time interacting with the novel mouse, whereas HSV-ZFP189^VPR^ treated mice do not. * p-value < 0.05, ns p-value > 0.05. n = 7 mice (HSV-ZFP189^DN^), n = 9 mice (HSV-ZFP189^WT^), n = 10 mice (HSV-ZFP189^VPR^). **(H)** Experimental timeline for tube test. For five days, social hierarchy was determined by once daily head-to-head tube tests for all animals within a five-mouse cage. In each cage, either the two most socially dominant or the two most subordinate mice were delivered HSV-ZFP189 TFs intra-PFC whereas the remaining cage-mates were delivered HSV-GFP. Mice recovered for one day. For the following five days the tube tests were repeated and wins for the HSV-ZFP189 TF treated mice versus all HSV-GFP treated mice was recorded. **(I)** Delivering HSV-ZFP189^VPR^ to the PFC of previously socially dominant mice decreases the probability of winning a tube test to a rate of random chance (dotted line). Two-way repeated-measures ANOVA comparing main effect of wins pre-vs. post-viral intervention. *** p-value < 0.0001. n = 4 cages with two HSV-ZFP189^VPR^ mice per cage. **(J)** Delivering HSV-ZFP189^VPR^ to the PFC of previously socially subordinate mice increases the number of wins per day to a 50% chance of winning against an opponent. Two-way repeated-measures ANOVA comparing main effect of wins pre-vs. post-viral intervention. ** p-value < 0.005. n = 6 cages with two HSV-ZFP189^VPR^ mice per cage. **(K)** Delivering HSV-ZFP189^DN^ to the PFC of socially dominant mice does not change their tube test wins from pre-surgery performance. Two-way repeated-measures ANOVA comparing main effect of wins pre-vs. post-viral intervention. ns p-value > 0.05. n = 4 cages with two HSV-ZFP189^DN^ mice per cage. **(L)** Delivering HSV-ZFP189^DN^ to the PFC of socially subordinate mice does not change their tube test wins from pre-surgery performance. Two-way repeated-measures ANOVA comparing main effect of wins pre-vs. post-viral intervention. ns p-value > 0.05. n = 6 cages with two HSV-ZFP189^DN^ mice per cage.

To understand the requirement of prior stress in driving these behaviors, and to more completely interrogate the nature of ZFP189^VPR^-induced social deficits, we performed three chamber social interaction testing in test mice with no prior stress experience and treated intra-PFC with our synthetic TFs (Fig. 3E). When provided the opportunity to interact with either a novel mouse or empty cage, mice expressing ZFP189^DN^ or ZFP189^WT^ show a normal preference for social interaction (Fig. 3F). Mice expressing ZFP189^VPR^ intra-PFC show no preference for social interaction over investigating an empty cage (Fig. 3F). When this test is repeated with the choice between the mouse from the previous round (familiar mouse) or a novel mouse, HSV-ZFP189^VPR^-treated test mice again show no ability to discriminate, or preference for, social interaction with the novel mouse over the familiar mouse (Fig. 3G). In performing these same studies in female mice, ZFP189^VPR^ identically ablates social behaviors (Supp. Fig. 3), revealing that ZFP189 regulates these social behaviors in both sexes. Collectively, these data further contextualize ZFP189^VPR^-driven social deficits to demonstrate that ZFP189^VPR^ behavioral control does not require prior stress experience, nor does it drive an active, anxiety-based social avoidance, as the ZFP189^VPR^-treated mice do not spend less time in the social chamber. Instead, ZFP189^VPR^ ablates an animal’s interest and/or awareness of social stimuli.

To understand the breadth of the detachment from social function induced by our synthetic ZFP189, we characterized the impact of ZFP189 dysregulation on an animal’s awareness and participation in prior established social hierarchy structures. We tested this by performing social dominance tube tests within each five-mouse cage before and after being manipulated with our synthetic ZFP189 TFs (Fig. 3H). Prior to viral-mediated gene transfer, we observe an establishment and maintenance of social hierarchy amongst cage-mates. In socially dominant mice, characterized as consistently winning a majority of the tube-tests against familiar cage-mates, viral delivery of ZFP189^VPR^ to the PFC ablates consistent wins in the tube test versus cage-mates and the chance of winning regresses towards 50%, which could be explained by random chance in the binary win/loss outcome of a tube test (Fig. 3I). Conversely, in socially subordinate mice, characterized as losing the most tube tests against cage-mates, viral delivery of ZFP189^VPR^ to the PFC increases the consistent wins in the tube towards winning half of the total bouts (Fig. 3J). Thus, both animals that were previously socially dominant or subordinate deviate from their previously established position in a social structure and regress to a social performance that could be explained by random chance. Critically, identically performed experiments with dominant and subordinate mice treated with ZFP189^DN^ do not show this deviation from their established social position (Fig. 3K-L), suggesting that this loss of hierarchical maintenance is specifically due to ZFP189^VPR^ PFC molecular function. This experiment indicates that ZFP189 dysfunction in the PFC removes the animal’s capacity to recognize and participate in cooperatively established social structures, like dominance hierarchies.

We next sought to interrogate if any of our observed behavioral deficits could be explained by ZFP189^VPR^-driven deficits in social memory. We performed a five-trial social memory test wherein mice manipulated with our synthetic ZFP189 TFs were subjected to four sequential sessions of social interaction with the same social target mouse and a fifth session wherein a novel mouse is introduced. Mice in all treatment groups significantly decreased time spent interacting with the increasingly familiar target mouse across sessions 1-4 and increased social interaction when the novel mouse was introduced (Supp. Fig. 4). As expected, animals manipulated with ZFP189^VPR^ interacted with the target mouse less overall (Supp. Fig. 4). These data indicate that ZFP189^VPR^ does not ablate the entirety of an animal’s social function, as these animals are still able to discriminate and remember novel vs. familiar conspecifics. Lastly, we sought to ensure that ZFP189^VPR^ does not cause deficits in inanimate object recognition. We performed a novel object recognition test and discovered that mice in all treatment groups were able to recognize novel objects (Supp. Fig. 5). These data indicate highlight the specificity of ZFP189^VPR^-induced behavioral deficits, in that it disrupts the higher-order social cognition necessary for group living, but not the more basic social and object recognition.

**Figure 4:**
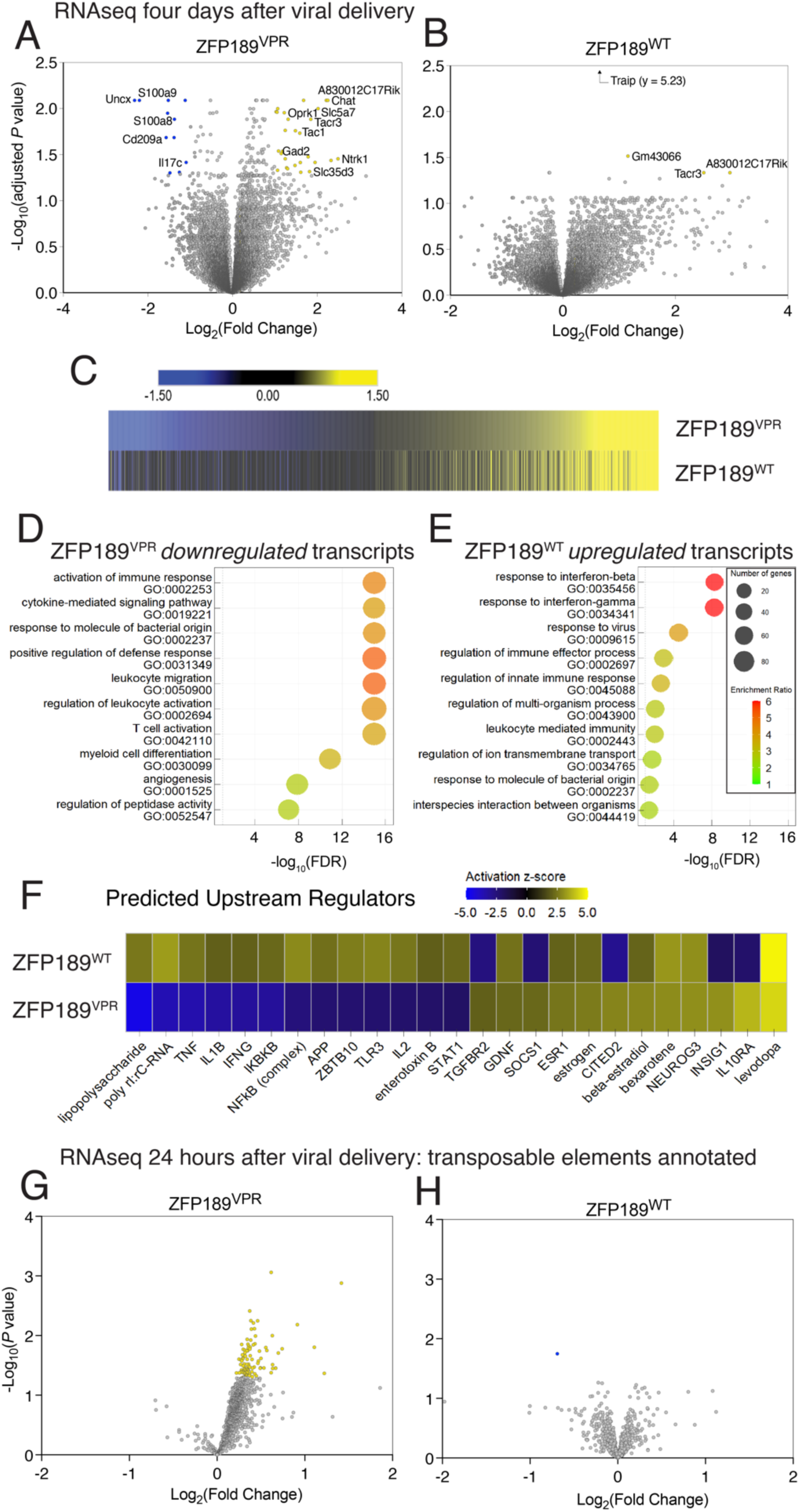
Synthetic ZFP189 TFs oppositely regulate transposable elements and genes related to immune function. **(A)** Four days after viral delivery, RNAseq was performed. Volcano plot depicting differentially expressed genes (DEGs) regulated by HSV-ZFP189^VPR^ in prefrontal cortex (PFC). Up-regulated genes are indicated by yellow dots, and down-regulated genes are indicated by blue dots. DEGs generated relative to our neutral HSV-ZFP189^DN^ comparator. DEG significance is set to 5% FDR corrected, Wald test adjusted *P* value < 0.05 and absolute log2 fold change > 1. n = 7 microdissected PFC from individual mice (HSV-ZFP189^DN^), n = 8 microdissected PFC from individual mice (HSV-ZFP189^VPR^). **(B)** Volcano plot depicting DEGs regulated by HSV-ZFP189^WT^ in PFC. DEGs generated relative to our neutral HSV-ZFP189^DN^ comparator. DEG significance as in (A). n = 7 microdissected PFC from individual mice (HSV-ZFP189^DN^), n = 5 microdissected PFC from individual mice (HSV-ZFP189^WT^). **(C)** Union heatmap comparing the pattern of HSV-ZFP189^VPR^ regulated transcripts organized from down-regulated to up-regulated (top) to HSV-ZFP189^WT^ transcripts of the same identity (bottom). Significance cutoffs for this and subsequent analyses nominal *P* value < 0.05 and absolute log2 fold change > 1.3, to enable broad pattern recognition. **(D-E)** Gene ontology (GO) biological function terms enriched within the DEGs uniquely down-regulated by ZFP189^VPR^ (D) and from the DEGs uniquely up-regulated by ZFP189^WT^ (E) reveal that both DEGs lists are comprised of DEGs whose gene products participate in immune-related biological functions. **(F)** Heatmap derived from Ingenuity pathway analysis (IPA) performed on complete DEGs lists reveals molecular factors whose function could explain the DEGs in our differential lists. HSV-ZFP189^WT^ vs. HSV-ZFP189^VPR^ regulated DEGs could largely be explained by increased (yellow) vs. decreased (blue) function of immune factors, respectively. **(G)** 24 hours after viral delivery, RNAseq was performed and transposable elements (TEs) were annotated in the reference library. In HSV-ZFP189^VPR^ manipulated PFC, hundreds of distinct TEs are up-regulated. TE DEGs generated relative to HSV-ZFP189^DN^. n = 5 microdissected PFC from individual mice (HSV-ZFP189^DN^), n = 5 microdissected PFC from individual mice (HSV-ZFP189^VPR^). **(H)** In HSV-ZFP189^WT^ manipulated PFC, only one TE is detected as down-regulated relative to the control HSV-ZFP189^DN^. n = 5 microdissected PFC from individual mice (HSV-ZFP189^DN^), n = 5 microdissected PFC from individual mice (HSV-ZFP189^WT^).

### ZFP189 regulates TEs and immune genes in prefrontal cortex

The gene regulatory functions of ZFP189 are poorly understood. To uncover the PFC transcripts regulated by ZFP189 that may be associated with these social deficits, we performed RNAseq in micro-dissected PFC virally over-expressing synthetic ZFP189 TFs. We generated differentially expressed genes (DEGs) relative to the control HSV-ZFP189^DN^ treatment four days post viral surgery, which is consistent with the social behavioral timeline in Figure 3. In mice treated with HSV-ZFP189^VPR^, we observe a symmetrical up-and down-regulation of PFC DEGs, represented in a volcano plot (Fig. 4A, full DEG list in Supp. Table 1). The absence of highly up-regulated DEGs was unexpected since our luciferase reporter data (Fig. 1C) suggests that ZFP189^VPR^ is a robust gene activator. This hints that at this time point multiple days after initial *trans*-gene ZFP189^VPR^ expression, the direct gene-activation effects of ZFP189^VPR^ have ameliorated, or that there are biological mechanisms to constrain the magnitude of ZFP189^VPR^-mediated gene activation in brain. Interestingly, HSV-ZFP189^WT^ produces fewer significant DEGs and only produces significantly up-regulated transcripts (Fig. 4B, full DEG list in Supp. Table 2). The lncRNA *A830012C17Rik* and the protein coding gene *Tacr3* were significantly up-regulated by both ZFP189 TFs, implying some converging function secondary to the demonstrated opposite direct gene regulation for these two TFs.

To further explore the similarities and differences in function of our opposing ZFP189 TFs, we compared the patterns of PFC transcriptional regulation by setting significance cut-offs to a nominal *p*<0.05 and a fold change threshold of > ±1.3 and aligning DEG identities in a union heat map (Fig. 4C). While minimal concordance was observed in the identities of DEGs down-regulated by ZFP189^VPR^ or ZFP189^WT^, there was an unexpected similar pattern of up-regulated DEGs by either ZFP189 TF (Fig. 4C; Supp. Fig. 6A-B). ZFP189^VPR^ drives far more down-regulated DEGs than ZFP189^WT^, with minimal overlap in gene identities. Whereas ZFP189^VPR^, and to a greater degree, ZFP189^WT^ both drive an increase in up-regulated DEGs with a sizeable overlap in commonly up-regulated DEGs (Hypergeometric test *p*-value < 0.0001) (Supp. Fig. 6B). These results are surprising, since the net consequence of transcriptional control of each ZFP189 TF is opposite to the direct gene regulatory control exerted by the TF. This implies that many of our detected DEGs are regulated downstream of the direct gene-regulation of ZFP189, likely via the up-(ZFP189^VPR^) or down-regulation (ZFP189^WT^) of some repressive intermediate mechanism.

To understand the biological functions of the genes uniquely impacted by each ZFP189 TF, we employed gene ontology (GO) analyses. Transcripts uniquely down-regulated by ZFP189^VPR^ are related to activation of an immune response (Fig. 4D). Similarly, transcripts up-regulated by ZFP189^WT^ are also immune-related, with particular emphasis on the interferon adaptive immune system (Fig. 4E). In the commonly up-regulated transcripts, we observe that both ZFP189^WT^ and ZFP189^VPR^ converge in activating the processes related to synaptic function, including neuropeptide and neurotransmitter signaling (Supp. Fig. 6C). This supplements our earlier observation that any form of ZFP189-mediated transcription function in PFC potentiates synaptic maturity on pyramidal neurons (Fig. 2).

To more completely interrogate the PFC gene networks regulated by ZFP189, we employed ingenuity pathway analysis (IPA) to identify and contextualize potential transcriptional regulators of our identified DEGs. Strikingly, ZFP189^VPR^ - vs. ZFP189^WT^ -regulated transcripts could be explained by increased (ZFP189^WT^) or decreased (ZFP189^VPR^) function of immune processes attributed to tumor necrosis factor (TNF), toll-like receptor 3 (TLR3), Interleukin 1β (IL1B), and/or Interferon γ (IFNG; IFN-ɣ), among others (Fig. 4F; Supp. Fig. 6D-E). This result builds on our GO analyses and further reinforces the divergent impact on immune processes achieved by our two distinct ZFP189 TFs, with ZFP189^WT^ activating, and ZFP189^VPR^ inhibiting, the PFC expression of these immune-related genes.

To illuminate the potential molecular mechanisms by which our distinct ZFP189 TFs achieve these unique patterns of immune gene expression, we sought to capture the DEGs most immediately regulated by our ZFP189 TFs. We performed RNAseq 24-hours after virally delivering ZFP189 TFs to the PFC. Given the major role that KZFPs like ZFP189 play in regulating the expression of TEs, we hypothesized a potential role for TEs in this mechanism. We modified RNAseq reference libraries to annotate TEs and generated differentially expressed TEs relative to the ZFP189^DN^ condition, as before. ZFP189^VPR^ activated the expression of more than 100 distinct TEs, and no TEs were detected to be down-regulated (Fig. 4G, differential TEs in Supp. Table 3). ZFP189^WT^ had a far more negligible effect on the expression of TEs relative to control conditions with only one down-regulated TE detected (Fig. 4H, differential TEs in Supp. Table 4). This ∼100-fold increase in the number of TEs transcribed by ZFP189^VPR^ indicates that ZFP189^VPR^ directly binds TE-rich regions of DNA and activates the expression of proximal TEs. The use of synthetic ZFP189 illuminates the natural sequence of molecular events, with ZFP189 directly binding TE-rich regions of DNA which consequently increases the expression of immune-related genes in PFC.

## Discussion

By designing synthetic ZFP189 TFs capable of exerting opposing forms of transcriptional control in the brain, we are able to identify that ZFP189-mediated transcriptional regulation in PFC governs social behaviors in rodents. By inverting the natural, repressive gene-regulation of ZFP189^WT^ to gene-activation with ZFP189^VPR^, we observe that mice demonstrate pronounced deficits in social cognition with no other detectable behavioral abnormalities. In combining our transcriptional manipulations with RNAseq transcriptome profiling, we detect that ZFP189^VPR^ consequently down-regulates, and ZFP189^WT^ up-regulates, immune genes that comprise the anti-pathogen response, suggesting that the PFC function of ZFP189 has evolved to regulate these genes as a means of connecting brain immune function to social behaviors.

Our data are consistent with growing evidence to support the co-evolution of brain immune functions and the maintenance of proper social behavior. Since group-based social behaviors are critical to the survival of many species, and aggregation of organisms increases the likelihood of spreading pathogens, there may have been a co-evolutionary pressure for anti-pathogen immune signaling cascades to also regulate the neurobiological processes that drive an organism’s capacity for group-based social behaviors. Earlier reports have indicated that the function of adaptive immune signaling in PFC, specifically IFN-ɣ, drives prosocial behaviors(12). It was demonstrated that IFN-ɣ deficient mice displayed social impairment without anxiety or motor deficits, which was reversible with administration of exogenous IFN-ɣ. Other research groups have uncovered that IL-17 signaling in neurons promotes social behaviors in mice (13). In our data, ZFP189^WT^ up-and ZFP189^VPR^ down-regulates the expression of genes whose products are involved in IFN-ɣ signaling cascades, and *Il17c* and IL-17 receptor genes are among the most down-regulated transcripts in the PFC of ZFP189^VPR^ treated mice (Fig. 4A). This reveals that, among other functions, ZFP189 regulates the transcription of genes involved in PFC immune signaling.

It is appreciated that a major evolutionary driving force for the expansion of the KZFP TF family is to control the expression of TEs (4), including in the adult brain (3,5), as a means of limiting their activity and potentially negative effects. We discover that ZFP189^VPR^ releases many TEs in the PFC (Fig. 4G), suggesting that ZFP189 normally binds and regulates TE-rich regions of the genome. There is a robust and growing literature to indicate that TEs have evolved to act as *cis-* regulatory elements(14), particularly at enhancer regions for interferon and immune-related genes (15,16). We hypothesize that the active transcription of these TE-rich gene loci, induced by our synthetic ZFP189^VPR^, incapacitates the *cis-*regulatory functions of these TE-rich DNA loci and inhibits the transcription of the predominantly immune-related *cis*-regulated genes. This potential mechanism would explain the gene-expression profiles we observe in our RNAseq studies, and is represented as a graphic in Supplemental Figure 7

Since HSVs specifically infect neurons (17), our synthetic ZFP189 TF over-expression is restricted to PFC nerve cells. However, the PFC cell-types that harbor our observed DEGs are not possible to ascertain given the bulk RNAseq approaches applied in this work. While IFN-ɣ is appreciated as originating from meningeal T cells, it is a soluble factor capable of acting upon IFN-ɣ receptors expressed on PFC microglia and neurons, with signaling in cortical neurons as solely implicated in regulating social behavior (12). Further, IL-17 signaling through neuronally expressed IL-17 receptors is required for social behaviors (13). Thus, we suspect that a majority of immune-related transcripts regulated by ZFP189 are localized within transduced PFC neurons, which would augment immune signaling within this cell type and consequently manifest the observed social deficits seen here.

Notably, we discovered a similar increase in mature spine density upon PFC pyramidal neurons facilitated by either ZFP189 TF. It is possible that these effects emerge as a product of ZFP189-mediated transcription functioning entirely within infected pyramidal nerve cells. This possibility is supported by the observation that ZFP189^VPR^ increases morphological complexity of N2a cells in a reduced cell culture preparation (Supp. Fig. 2). However, we also observed a converging effect on up-regulation of transcripts that are unique to, and drive the function of, PFC cholinergic neurons. Specifically, many of the top co-up-regulated DEGs such as *Lhx8, Ntrk1, Isl1, Chat, Six3, Nkx2-1, Slc5a7,* among others, encode gene products required for the development and function of PFC cholinergic neurons (18–20), suggesting that synaptic changes on pyramidal neurons may, in part, be the product of local functions of cholinergic interneurons, as has been previously reported (21,22). A small fraction of cortical neurons are the vasoactive intestinal polypeptide (VIP) and choline acetyltransferase (ChAT) positive cholinergic interneurons (23), which are identified by the unique expression these transcripts (24), and have been noted to increase excitatory neurotransmission upon PFC pyramidal neurons to control attention (21). This raises the interesting possibility that ZFP189-induced altered function of these VIP+/ChAT+ interneurons contributes to the lack of social attention observed in our HSV-ZFP189^VPR^ treated mice. Further, cholinergic neurotransmission in the forebrain has been documented as being both sensitive to and contributing to immune response (25–27). Still, it remains unclear if the morphological changes of PFC pyramidal neurons and the expression of VIP+/ChAT+ genes are a direct consequence of ZFP189-mediated regulation of immune genes, or a result of independent ZFP189 gene-regulatory functions, which are perhaps distinct within infected VIP+/ChAT+ interneurons. This is an important area for future studies.

We elected to employ our functionally inert ZFP189^DN^ TF as a neutral comparator control group in these studies since we established that it exerts no regulatory control at a luciferase ZFP189 target gene (Fig. 1C) and therefore most appropriately controls for off-target protein interactions as a result of viral over-expression of our synthetic TFs. Furthermore, we are confident that our observed effects in transcription of immune-related genes is not a consequence of viral mediated gene transfer itself for a number of reasons. First, in all of our RNAseq analyses, we generated DEGs against the HSV-ZFP189^DN^ condition, which is matched in terms of delivered viral particles and would therefore normalize out genes that respond to HSV infection *per se*. Second, many other studies have utilized identical viral-mediated gene transfer strategies, yet not observed an immune response in their transcriptomic analyses(28–30). Most importantly, immune-related genes are bi-directionally regulated depending on the delivered ZFP189 TF, and are therefore unlikely to be a response to viral delivery itself. This supports the interpretation that our virally delivered synthetic ZFP189 TFs are themselves responsible for regulating our observed DEGs.

Our data also complement a growing body of research that has linked the brain expression and function of specific TFs, such as EGR1, to social behaviors in many organisms including songbirds (31,32), fish (33), and rodents(34). Importantly, EGR1 (sometimes called NGFI-A, KROX24, or ZIF268) is a zinc finger TF, whose brain expression has been noted to respond to social experiences and drive pro-social behaviors across phylogeny (31–34). Given that ZFP189 is a member of a similar zinc finger TF gene family, it is possible that other structurally similar TFs, such as other KZFP TFs, regulate social behaviors via similar mechanisms of TE-regulated immune response. In support of this notion, the blind mole rat and the naked mole rat, two related organisms that differ widely in degree of TE regulation and immune responses (35–37) also represent extremes of social behaviors, ranging from highly solitary, in the case of the blind mole rat (38), to one of the few eusocial mammals with extremely complex and cooperative social structures, in the case of the naked mole rat (39). Also, higher social status amongst macaques drives a proinflammatory response (40) via chromatin reorganization to expose TF DNA motifs (41). Here, we observed that inverting ZFP189 gene-regulatory function in PFC removed the animal’s capacity to participate in cooperatively established group social structures, like social dominance hierarchies (Fig. 3I-J). This raises the intriguing possibility that eukaryotic brain TFs have co-opted the molecular mechanisms originally exploited by pathogens to influence social behaviors to facilitate pathogen transmission. It is possible that ZFP189 tunes the neurobiological mechanisms that facilitate social aggregation and group-based behaviors by regulating TE-rich regions of DNA, much of which is derived from ancient retroviruses(42), as a way of augmenting brain immune response and associated social behaviors, completely in the absence of external pathogens.

Lastly, ZFP189 was identified in an open-ended co-expression network analysis of RNAseq data across multiple limbic brain regions of mice that demonstrated phenotypic resilience to manifesting social deficits in response to chronic social stress (6). *Zfp189* was the top key driver gene within the resilient-specific gene-network and *Zfp189* viral overexpression or CRISPR-targeted gene activation in PFC was sufficient to manifest the resilient-specific gene network and endow the animal with behavioral resilience to chronic stress, as measured in social interaction tests as well as other behavioral endpoints that do not involve social behaviors (6). However, there was no indication that ZFP189 regulated the genes within the resilient-specific network via direct TF:gene interactions. Importantly, in post-mortem tissue from human PFC (Brodmann area 25), individuals with major depression had lower expression of *ZNF189* mRNA than matched controls. This raises the striking possibility that *Zfp189* expression in the PFC is biologically variable across individuals and sensitive to the experiences of the individual, such as the experience of chronic stress. This could mean that stress-induced social deficits observed in a number of neuropsychiatric syndromes are, in part, driven by PFC ZFP189 dysfunction and the TE-associated regulation of immune functions. This research is a critical area for future investigation.

Here we present a novel approach in re-programming the functions of a poorly understood TF. Only by synthetically inverting the gene-regulatory function of ZFP189 in the brain were we able to discover that ZFP189 coordinates complex social behaviors via regulation of genomic TEs and subsequent immune response in PFC. This work uncovers a novel connection between brain TF regulation of genetic transposons, immune response, and the social cognition necessary for group living.

## Acknowledgements

This work was supported by the NIH grants R00DA045795 and P30DA033934 to PJH. Microscopy was performed at the VCU Microscopy Facility, supported, in part, by funding from NIH-NCI Cancer Center Support Grant P30CA016059.

## Methods

### Subjects

Male and female C57BL/6J mice (8 to 10 weeks old) from Jackson Laboratories were used. Mice were group housed (5 mice/cage) on a 12 hour light/dark cycle (lights on at 6am/off at 6pm) with food and water freely available. All mice were used in accordance with protocols approved by the Institutional Care and Use Committees at Virginia Commonwealth University School of Medicine.

### Viral packaging

We *de novo* synthesized ZFP189^DN^, ZFP189^WT^, and ZFP189^VPR^ and sub-cloned into HSV expression plasmids via ThermoFisher Scientific gateway LR Clonase II cloning reaction and Gateway LR Clonase II Enzyme mix kit (catalog number 11791-020 and 11971-100). Colonies were Maxiprepped (Qiagen Cat # 12163) and shipped to the Gene Delivery Technology Core at Massachusetts General Hospital for HSV packaging. Once packaged, aliquots were made and stored in −80 degrees C to be used in viral gene transfer through stereotaxic surgery.

### Viral gene transfer

Stereotaxic surgeries targeting the PFC were performed as previously described (6,43). Mice were anesthetized with I.P. injection of ketamine (100 mg/kg) and xylazine (10 mg/kg) dissolved in sterile saline solution. Mice were then placed in a small-animal stereotaxic device (Kopf Instruments) and the skull surface was exposed. 33-gauge needles (Hamilton) were utilized to infuse 1.0 μL of virus at a rate of 0.2 μL/minute followed by a 5-min rest period to prevent backflow. The following coordinates were used to target the PFC: Bregma: anterior-posterior: +1.8mm, medial-lateral +0.75mm, dorsal-ventral −2.7mm, 15 degree angle (6).

### Neuro-2a Cell Culture and Transfection

*Mus musculus* Neuro-2a (N2a; ATCC^®^ CCL-131™) neuroblast cells were grown in adherent culture with EMEM growth medium (Quality Biological, #112-039-101; ATCC, #30 2003 or Corning, #10-009-CV) with 5% FBS (HyClone, #SH30071.03IH30-45) and 1.5% Penicillin Streptomycin (Gibco, #15140122) in a 37°C and 5% in a Thermo Scientific HERAcell-150i CO_2_ incubator using aseptic techniques. N2a cells were maintained by passaging twice per week. One day before transfection, N2a cells of passage 60 or fewer were seeded in a 96-well plate at approximately 1.5×10^4^ cells/well. On the following day, the plasmid DNA of RE reporter (25mg) was co-transfected on 80% confluent wells in replicate for all treatment conditions with QIAGEN Effectene Transfection Kit (#301427) according to manufacturer instructions. Each reaction was carried out in 10ul of buffer EC, 0.3ul of Enhancer and 1ul of Effectene, that was diluted to 100ul in the medium before applying to the cells. The corresponding empty reporter vector RL (LightSwitch, #S990005) and empty expression vector GFP (p1005gw Δ*CCDB*) were used as background controls. The plates were centrifuged for 7 minutes at 2000rpm for higher transfection efficiency and were incubated for 1-3 days with coving by Breathe-Easy sealing membrane only (Sigma-Aldrich, # Z380059-1PAK). All assays were performed on Day 1-3 following transfection. In experiments, each n corresponds to a distinct well, and experiments were replicated across 2-3 transfection reactions performed on different days.

### Luciferase assay

To assess the cell viability and toxicity, on day 2 following GFP reading, we conducted a Luciferase Assay Report using the CellTiter-Glo 2.0 Assay (Promega #G9242) and Renilla luciferase assay system (Promega #E2820). Relative Luminesce Unit (RLU) was measured by the BMG-Omega plate reader using the Luminescence endpoint program with gain 3432.

### N2a Morphology Assessment

The plasmid DNA of viral expression vectors (0.1-0.3ug) were transfected into N2a cells as described above in triplicate. The cell images of 48 hours were captured in 0.1ug transfection by a Bio-Rad ZOE_TM_. Fluorescent Cell Imager, followed by the GFP fluorescence intensity readout. For quantitative analysis of N2a morphology, we used Image J (FIJI) software to quantitate the number and length of each protrusion. We used the free-hand line tool to trace each protrusion with the set scale of 0.71pixel/um corresponding to the 100um scale bar on each Fluorescent Microscopy photo. We then used the ROI manager measure of lengths to calculate averages.

To further analyze our protrusion count and average protrusion length data, data from triplicate transfection reactions were averaged and normalized with the GFP readout.

### Microdefeat Stress

CD1 retired breeder mice were screened for aggressive behavior as previously described(44). A C57BL/6J mouse was subjected to subthreshold levels of social defeat that consist of three 5-min defeat sessions given consecutively on a single day with 15 min of rest between each session. In each defeat session, a C57BL/6J mouse is placed in the home cage of a single-housed CD1 mouse (44).

### Social Interaction (SI)

Behavioral testing was performed 24 hours after exposure to sub-threshold microdefeat stress. Social interaction was performed as previously described (44). C57BL/6J mice were placed into the open arena (43×43×43cm) with an empty wire cage (10×5×30cm) at one side (interaction zone). Mice were given 2.5 minutes of habituation to explore the arena and then removed from the open arena. A unfamiliar, novel CD1 aggressor was then placed within the wire cage (placed in interaction zone) and the C57BL/6J was placed back into the open arena and another 2.5 minutes were recorded. The arena was cleaned with 70% EtOH before a new C57BL/6J mouse was placed in the arena and before the first trial. Data were analyzed as time spent in the interaction zone without the CD1 target mouse compared to time spent within the interaction zone with the CD1 target present. The data was then calculated into SI ratios by dividing the time spent in the interaction zone with the target mouse present by the time spent in the interaction zone (IZ) with the target mouse absent (SI ratio = time in IZ target present / time in IZ target absent). Total distance moved (locomotion) was recorded and evaluated when the target was absent. Behavioral analyses were performed automatically by video tracking software (Ethovision Noldus) (45). All behavioral tests were performed in a specified behavioral suite under red light illumination.

### Elevated Plus Maze (EPM)

The EPM apparatus consists of two open arms (33×6cm) and two closed arms (33×9.5× 20cm) facing connected by a central platform (5×7cm) (46). The maze is elevated 63cm above the floor. A C57/BL6J mouse was placed individually in the right-side closed arm facing the center of the plus-maze. Placement of all four paws into an arm was registered as an entry in the respective arm. The time spent in each arm was recorded during the 5 min EPM test. The platform of the maze was cleaned with 70% EtOH following each trial and before the first trial. The percentage of time spent in the open arms was calculated (time spent in open arms/300 s * 100 = % time spent in open arms.) Behavioral analyses were performed automatically by video tracking software (Ethovision Noldus) (45). All behavioral tests were performed in a specified behavioral suite under red light illumination.

### Three Chamber Sociability and Social Novelty Test

The three-chamber test was used to assess sociability and interest in social novelty or social discrimination. The testing arena consisted of three adjacent chambers (each 41×21×41cm) separated by two clear plastic dividers and connected by open doorways (5×9cm). The test consisted of three 10-min sessions. The subject mouse begins the session in the middle chamber. In the first session, subject mice were allowed to habituate to the arena and freely investigate the three chambers. In the subsequent sociability session, a novel C57BL/6J same-sex mouse (target mouse 1) was placed in a cylindrical cage (20cm height×10 cm diameter solid bottom; with clear bars spaced 2 cm apart) in one of the side chambers and another identical inverted empty cup was placed in the other side chamber. In the social novelty session, the empty pencil cup was removed and replaced by target mouse 1 in a new pencil cup. A second novel C57BL/6J same-sex mouse (target mouse 2) was placed at the previous position of target mouse 1 under a new pencil cup. Target mice were the same sex and age as the subject mice. The chamber and cages cups were cleaned with 70% EtOH between animals and before the first animal. Time spent in each chamber was recorded. Sociability was measured by comparing the time spent in the chamber with a novel mouse vs an empty cup. Social novelty was measured by comparing the time spent in the chamber with a novel vs familiar mouse.

Behavioral analyses were performed automatically by video tracking software (Ethovision Noldus) (45) All behavioral tests were performed in a specified behavioral suite under red light.

### Social Dominance Tube Test

Animal social dominance was tested as previously described (47,48) in a transparent Plexiglas tube measuring 30.5cm in length and 3cm diameter, a size just sufficient to permit one subject mouse to pass through without reversing direction. The tube was set on a plastic table in the designated behavioral suite and trials were manually recorded by a researcher blind to experimental groups. Animals were placed at opposite ends of the tube and released. A subject was declared the “winner” when its opponent backed out of the tube, with all 4 paws outside of the tube. The maximum test time allowed was 2 minutes. For five days, baseline social hierarchy was determined by once-daily tube tests for all animals within a five-mouse cage in a randomized order. In each cage, either the two most socially dominant or the two most subordinate mice were delivered HSV-ZFP189 TFs whereas the remaining cage-mates were delivered HSV-GFP. For the following five days the tube tests were repeated and wins for the HSV-ZFP189 TF treated mice versus all HSV-GFP treated mice was recorded to determine post-surgery social hierarchy. Social dominance was measured by calculating the percentage of wins in the tube test (number of wins/number of tests * 100).

### Five-Trial Social Memory Test

Five-trial social memory was tested as previously described (49) to determine ability to recognize novel versus familiar animals. Subjects were placed into the open arena (43×43×43cm) with an empty wire cage (10×5×30cm) at one side (interaction zone). Mice were given 2.5 minutes of habituation to explore the arena and then removed from the open arena. A novel C57BL/6J male mouse was then placed within the wire cage (interaction zone) and the C57BL/6J was placed back into the open arena and another 2.5 minutes were recorded (trial 1). The novel mouse was removed for 10 minutes. Subsequently, the same procedure was repeated three more times (i.e. subject mouse exposed to the familiar mouse, trials 2, 3 and 4). In trial 5, an unfamiliar male C57BL/6J mouse was introduced to measure dishabituation. Time spent in the interaction zone for the first 30 seconds of each trial was measured. Behavioral analyses were performed automatically by video tracking software (Ethovision Noldus) (45). All behavioral tests were performed in a specified behavioral suite under red light.

### Novel Object Recognition Test

Novel object recognition was tested as previously described (50) In the first trial, the C57BL/6J mouse was placed in the center of an open arena (43×43×43cm) with two identical objects on opposite sides of the box, and was allowed to freely explore the objects for five minutes. In the second trial, (the object recognition memory testing phase) the mouse was allowed to explore objects for 5 minutes in the same open field box compromising one object used in the trial 2 (familiar object) and a new object replacing the second object used in trial 1 (the novel object.) A mouse is considered to be exploring an object when its nose is within 3 cm of the object. The arena was cleaned with 70% EtOH before a new C57BL/6J mouse was placed in the arena and before the first trial. Times spent exploring the novel object, the familiar object, and both objects (total object exploration time) were collected. Furthermore, novel object discrimination index was calculated (time spend with novel object/total exploration time * 100.) Behavioral analyses were performed automatically by video tracking software (Ethovision Noldus) (45). All behavioral tests were performed in a specified behavioral suite under red light.

### Microscopy

Three days following viral gene transfer, mice were transcardially perfused with 0.1 M sodium phosphate buffer, followed by 4% paraformaldehyde (PFA) in 0.1 M phosphate buffer. Brains were removed and postfixed in 4% PFA overnight at 4°C. Following postfix, whole brains were stored in 15% sucrose in phosphate buffer with 0.05% sodium azide at 4°C for 24 hours, and then stored in 30% sucrose in phosphate buffer with 0.05% sodium azide at 4°C until sectioning. Coronal sections at 40μm thick containing the PFC were then prepared on a Lecia VT1000S vibratome in 0.1 PBS. Sections were then mounted and cover slipped using ProLong Gold Antifade. Slides were kept at 4°C until imaging in a light-blocking slide box until imaging.

Sections were imaged and captured with Zeiss 880 Airyscan confocal microscope. Slices were imaged with a 63x oil-immersion magnification.

### Dendritic Spine Analysis

Images were captured on a Zeiss 880 Airyscan confocal microscope, and individual dendritic segments were focused on and scanned at 0.69-μm intervals along the z axis to obtain a z-stack. After capture, all images were deconvolved within the Zeiss Application Suite software. Analyses were performed on two-dimensional projection images using ImageJ (NIH). ∼20 μm in length of dendrite were analyzed on captured neurons. For each group, 3-6 cells were analyzed in 3-5 different mice. We operationally divided spines into three categories; (1) mushroom-like spines were dendritic protrusions with a head diameter >0.5μm or >2x the spine neck diameter; (2) stubby spines were dendritic protrusions with no discernable head and a length of ≤0.5μm; and (3) thin/filopodia-like spines were dendritic protrusions with a length of >0.5 μmand head diameter <0.5μm or no discernable head (51).

### Tissue Preparation and RNA Sequencing

Mice were virally delivered HSV-ZFP189^DN^, −ZFP189^WT^, or -ZFP189^VPR^ intra-PFC to be used in RNAseq analysis. Mice were cervically dislocated and decapitated without anesthesia, and the brains were removed and sectioned into 1 mm coronal slices using brain matrices. Bilateral tissue punches from the PFC (12 gauge; internal diameter, 2.16 mm) were snap frozen on dry ice and stored at −80°C, as is routinely performed by our group(52,53). Total RNA was extracted from fresh frozen tissue samples using Qiagen RNeasy Plus Universal mini kit following manufacturer’s instructions (Qiagen, Hilden, Germany). RNA samples were quantified using Qubit 2.0 Fluorometer (Life Technologies, Carlsbad, CA, USA) and RNA integrity was measured using the RNA Screen Tape on Agilent 2200 TapeStation (Agilent Technologies, Palo Alto, CA, USA). The average RNA integrity number for all samples exceeded 8.6. Samples were initially treated with TURBO DNase (Thermo Fisher Scientific, Waltham, MA, USA) to remove DNA contaminants. The next steps included performing rRNA depletion using QIAseq® FastSelectTM−rRNA HMR kit (Qiagen, Germantown, MD, USA), which was conducted following the manufacturer’s protocol. RNA sequencing libraries were constructed with the NEBNext Ultra II RNA Library Preparation Kit for Illumina by following the manufacturer’s recommendations.

Briefly, enriched RNAs are fragmented for 15 minutes at 94°C. First strand and second strand cDNA are subsequently synthesized. cDNA fragments are end repaired and adenylated at 3’ends, and universal adapters are ligated to cDNA fragments, followed by index addition and library enrichment with limited cycle PCR. Sequencing libraries were validated using the Agilent Tapestation 4200 (Agilent Technologies, Palo Alto, CA, USA), and quantified using Qubit 2.0 Fluorometer (ThermoFisher Scientific, Waltham, MA, USA) as well as by quantitative PCR (KAPA Biosystems, Wilmington, MA, USA). The sequencing libraries were multiplexed and clustered on one lane of a flowcell. After clustering, the flowcell was loaded on the Illumina HiSeq 4000 instrument according to manufacturer’s instructions. The samples were sequenced using a 2×150 Pair-End (PE) configuration at a sequencing depth of approximately 101 million reads per sample (mean = 101 ± 23 million). Other overall sample sequencing statistics include the mean quality score (37.50 ± 0.01) and the percent of bases ≥30 (94.76 ± 0.02).

Sequence reads were trimmed to remove possible adapter sequences and nucleotides with poor quality using Trimmomatic v.0.36. The trimmed reads were mapped to the Mus musculus GRCm38 reference genome available on Ensembl using the STAR aligner v.2.5.2b. Unique gene hit counts were calculated by using featureCounts from the Subread package v.1.5.2. For Figure 4G-H, transposable element hit counts were calculated using featureCounts from the TEtranscripts package(54) and were calculated alongside canonical DEGs. After extraction of gene hit counts, the gene hit counts table was used for downstream differential expression analysis. Using DESeq2, a comparison of gene expression between groups of samples was performed. Volcano plots with these DEGs were assembled using Graphpad Prism 9.

### DEG analysis approach

DEG tables from DESeq2 were imported into Rstudio for further analysis. For pattern identification in union heatmaps, Venn diagrams, and Gene Ontology (GO), genes with a nominal *p* value < .05 and absolute log_2_ fold change > 1.3 were defined as DEGs for each comparison. In preparation for union heatmaps, DEG lists were sorted by increasing log_2_(FoldChange). For Venn diagrams, the significant DEGs were segregated by up-or downregulation by the sign of log_2_(FoldChange), and the significance of overlap was calculated using a hypergeometric test. Both union heatmaps and Venn diagrams were constructed using the ggplot package (H. Wickham. ggplot2: Elegant Graphics for Data Analysis. Springer-Verlag New York, 2016). WebGestalt (55) was used for GO over-representation analysis on Common upregulated, WT upregulated, and VPR downregulated gene lists (Benjamini-Hochberg adjusted false discovery rate < 0.05). Outputs were graphically represented by fold enrichment and number of genes in each ontology per a script from Bonnot et al (10.21769/BioProtoc.3429). Briefly, enriched ontology terms were ordered by false discovery rate (FDR) and plotted using ggplot in Rstudio, with each plot point colored by enrichment and sized by number of genes. Finally, DEG lists were analyzed using Ingenuity Pathway Analysis to determine upstream regulators(56). Significantly activated and inhibited upstream regulators were sorted and graphed into an activation heatmap using the ggplot package.

---

**Supplemental Figure 1:**
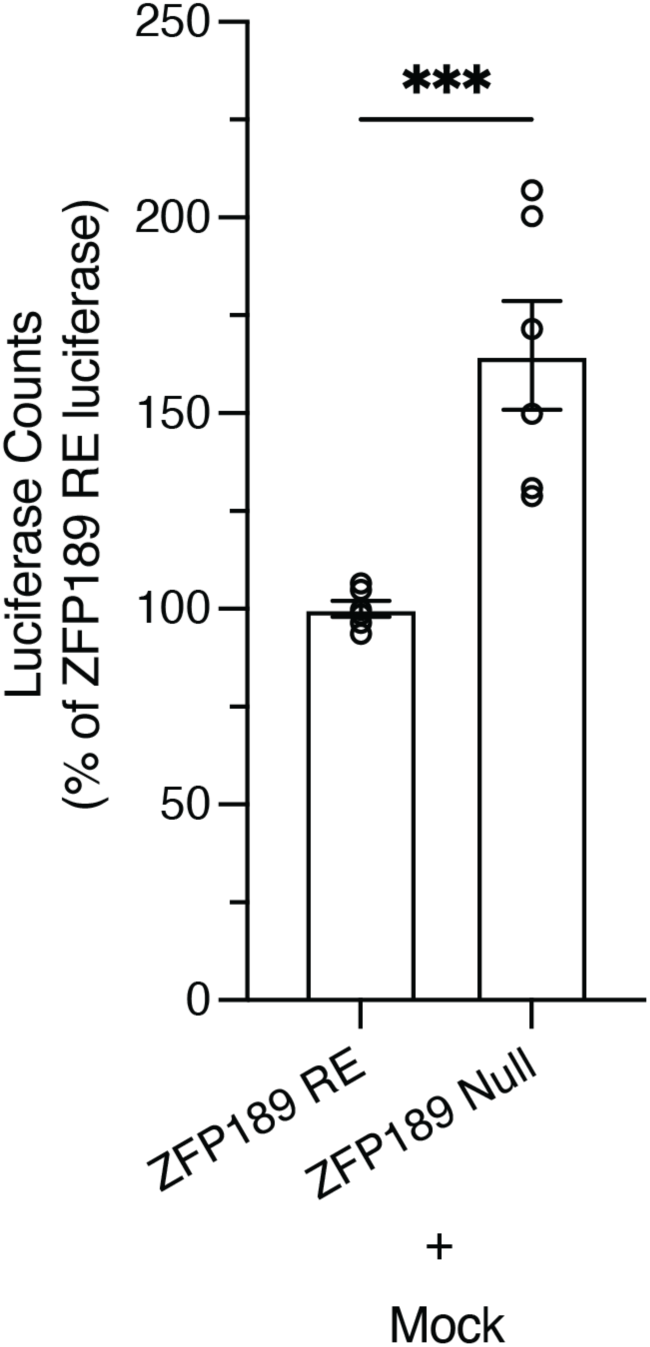
Transfecting ZFP189 luciferase reporters either possessing or lacking the ZFP189 DNA Response Element (RE) reveal the natural gene-repressive functions of ZFP189 endogenously expressed in N2a cells. N2a cells transfected with luciferase reporter target plasmids possessing (ZFP189 RE) or lacking (ZFP189 null) the ZFP189 RE promoter sequence uncovers the endogenous repressive action of ZFP189 in the mouse neuronal N2a cells. Removing ZFP189 DNA binding domains in the ZFP189 null condition increases luciferase expression, revealing a release of a repressive effect mediated by an N2a endogenously expressed protein that binds ZFP189 REs. Data normalized to ZFP189 RE condition, by experiment. Two-tailed, unpaired Student’s *t*-test; **** p-value < 0.001. n = 6 per condition.

**Supplemental Figure 2:**
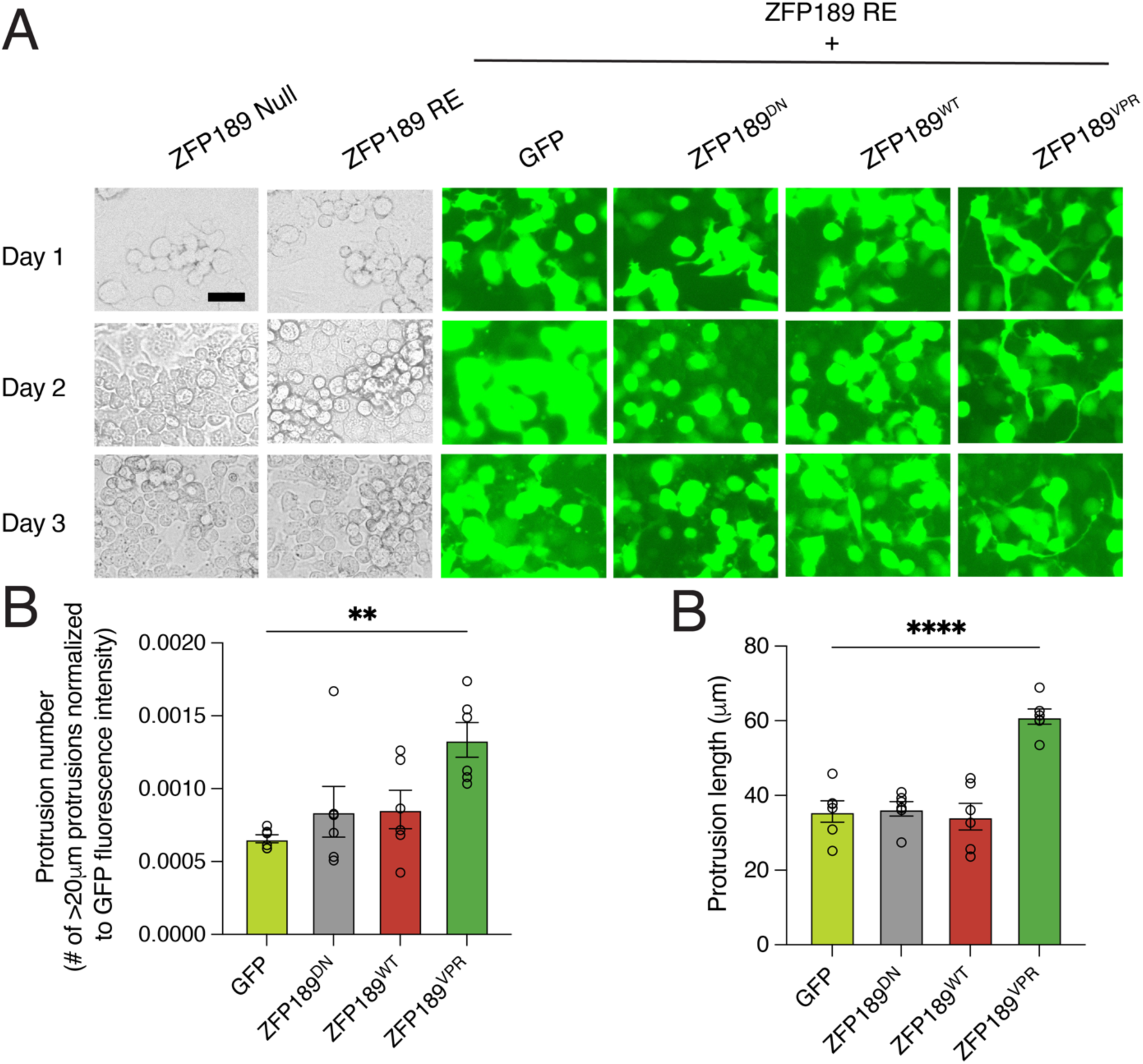
Transfecting N2a cells with ZFP189^VPR^ increases morphological complexity by increasing the number and length of cellular protrusions. **(A)** Microscopy on N2a cells transfected with only luciferase reporter plasmids (phase contrast; no plasmid GFP cassette in luciferase plasmids) or ZFP189 RE luciferase plasmid alongside synthetic ZFP189 TFs (fluorescent microscopy; GFP driven by separate CMV promoter in TF plasmids) for three days following the transfection reaction. Scale bar is 20 μm. **(B)** N2a cells expressing ZFP189^VPR^ have more cellular protrusions. On day 2, in each GFP+ condition, the total number of cellular protrusions in GFP+ cells measuring greater than 20 μm was quantified in each well and normalized to GFP fluorescence intensity for that well, to account for transfection efficiency. One-way ANOVA followed by Bonferroni’s multiple comparisons test relative to GFP condition. ** p-value < 0.01; n = 6 per condition. **(C)** N2a cells expressing ZFP189^VPR^ have longer cellular protrusions. On day 2, the length of each cellular outgrowth was measured and averaged per well. One-way ANOVA followed by Bonferroni’s multiple comparisons test relative to GFP condition. **** p-value < 0.0001; n = 6 per condition.

**Supplemental Figure 3:**
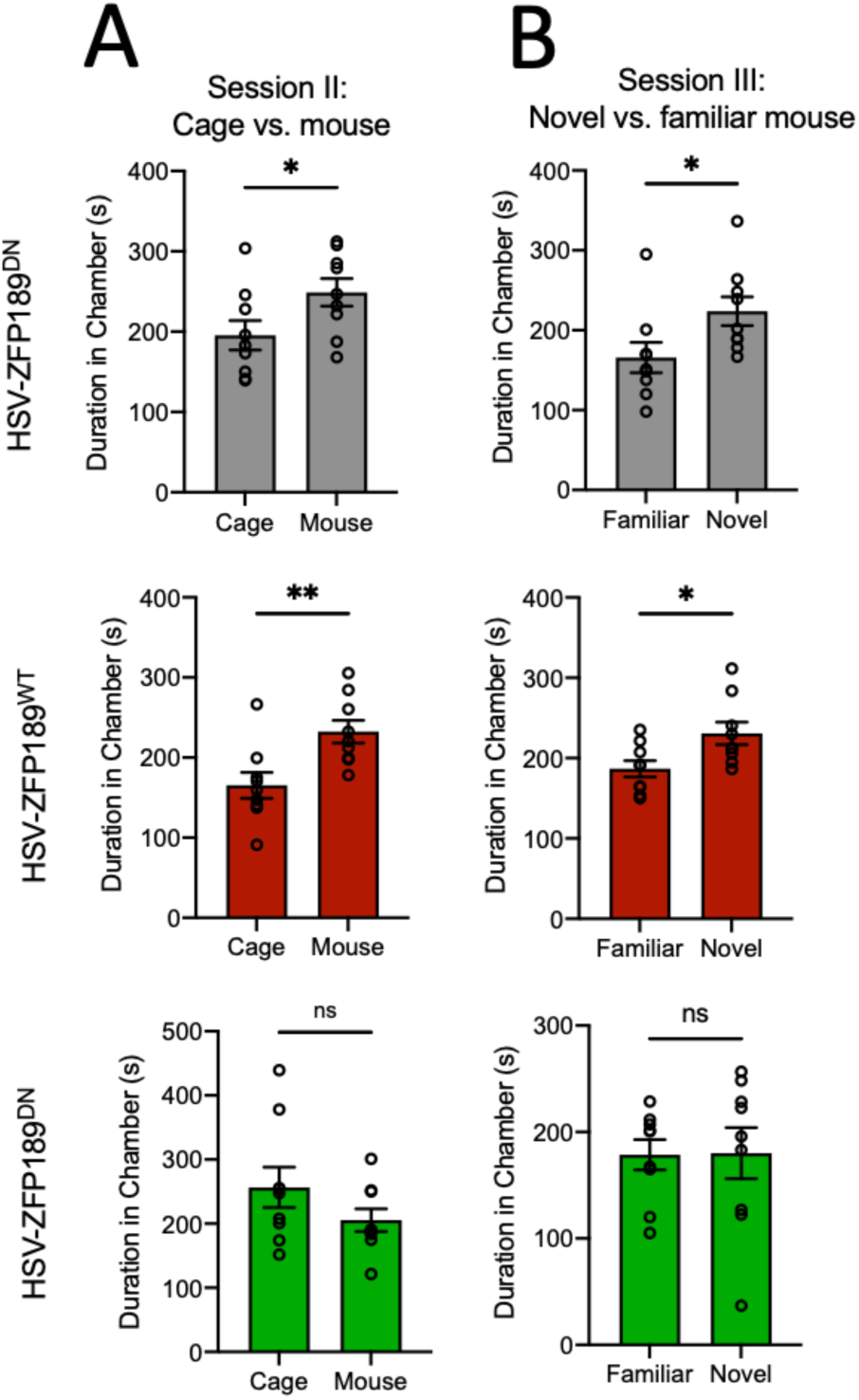
Inverting the natural transcriptional control of ZFP189 in the PFC with HSV-ZFP189^VPR^ impacts social function in female mice. **(A)** In session II, when given the choice between interacting with an empty cage or a novel mouse, HSV-ZFP189^DN^ and HSV-ZFP189^WT^ treated female mice spend more time interacting with the novel mouse, whereas HSV-ZFP189^VPR^ treated mice do not. Unpaired two-tailed Student’s *t*-test. ** p-value < 0.005, * p-value < 0.05, ns p-value > 0.05. **(B)** Subjecting these same animals to session III wherein female test mice are given a choice between interacting with the caged mouse from the previous round (Familiar) or a novel mouse, HSV-ZFP189^DN^ and HSV-ZFP189^WT^ treated mice again spend more time interacting with the novel mouse, whereas HSV-ZFP189^VPR^ treated mice do not. * p-value < 0.05, ns p-value > 0.05. n = 9 mice (HSV-ZFP189^DN^), n = 9 mice (HSV-ZFP189^WT^), n = 9 mice (HSV-ZFP189^VPR^).

**Supplemental Figure 4:**
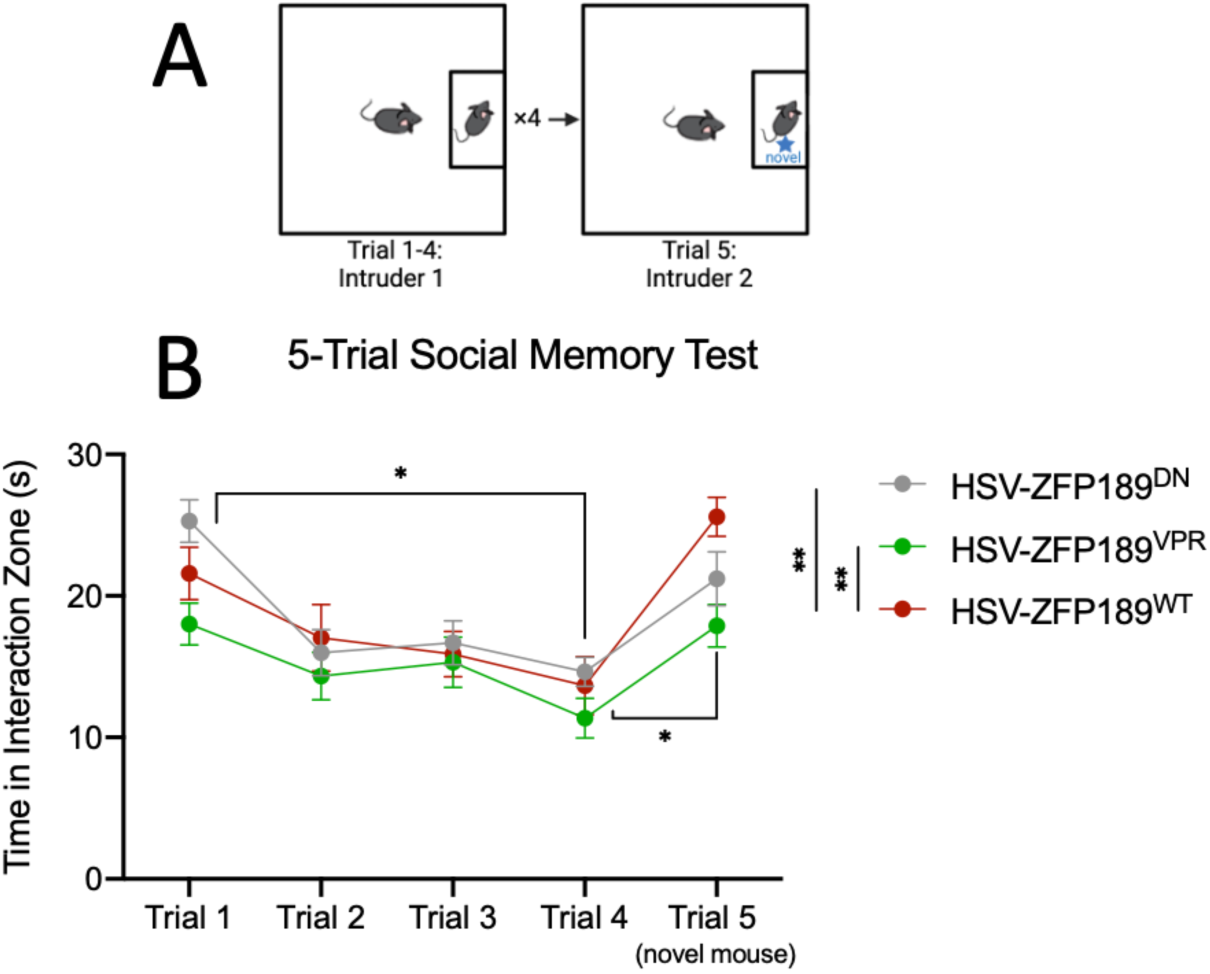
No viral treatment condition impacts social recognition or social memory. **(A)** Schematic of five-trial social memory paradigm. Mice manipulated with our synthetic ZFP189 TFs were placed in an arena with an unfamiliar target mouse for a 2.5-min trial duration. This is repeated with the same test mouse and target mouse for trials 1 through 4. In the 5^th^ trial, a novel mouse was introduced as the target mouse. Total time interacting with target mouse during the first 30 seconds of the trial is shown for each trial. **(B)** Mice in all groups show habituation to the familiarity of the test mouse in trials 1-4. All mice increase time spent socializing with the novel mouse in trial 5, indicating normal social memory in all groups. Tukey’s multiple comparisons test. 1 vs. 4, **** p-value < 0.0001; 4 vs. 5, * p-value < 0.05 (HSV-ZFP189^DN^), 1 vs. 4, * p-value < 0.05; 4 vs. 5, **** p-value < 0.0001 (HSV-ZFP189^WT^), 1 vs. 4, * p-value < 0.05; 4 vs. 5, * p-value < 0.05 (HSV-ZFP189^VPR^). Additionally, over all tests, HSV-ZFP189^VPR^ treated mice interacted with social target mice less than HSV-ZFP189^DN^ and HSV-ZFP189^WT^ treated mice. Tukey’s multiple comparisons test. ** p-value < 0.005. n = 14 mice (HSV-ZFP189^DN^), n = 12 mice (HSV-ZFP189^WT^), n = 15 mice (HSV-ZFP189^VPR^).

**Supplemental Figure 5:**
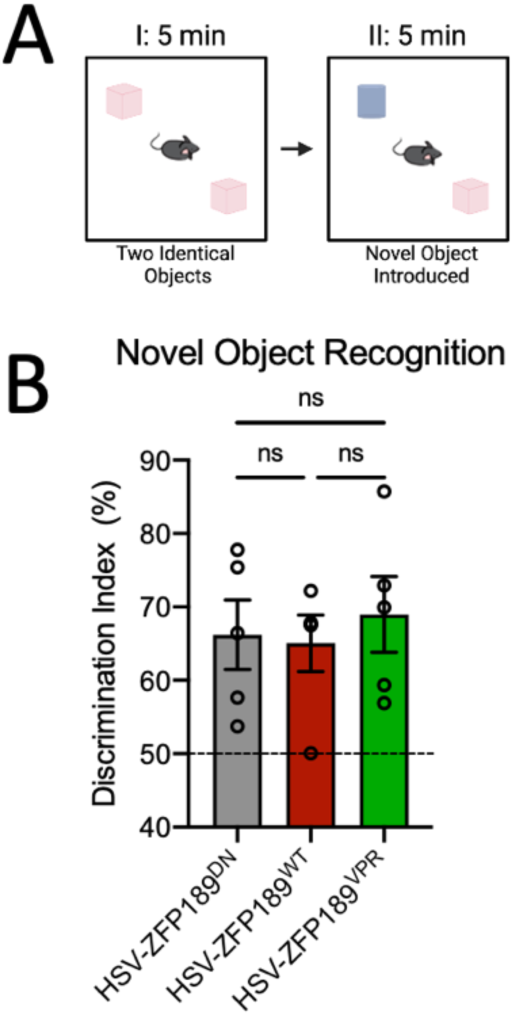
No viral treatment condition impairs novel object recognition. **(A)** Schematic showing novel object recognition paradigm. Mice manipulated with our synthetic ZFP189 TFs were placed in arena with two identical objects on opposite sides of the box, and were allowed to freely explore the objects for five minutes. In the second trial, the mouse was allowed to explore objects for 5 minutes in the same arena compromising one object used in the trial 2 (familiar object) and a new object replacing the second object used in trial 1 (the novel object.) **(B)** Represents the discrimination index percentage (DI) for each experimental group. DI = time spend with novel object/total exploration time * 100. All experimental groups were able to recognize novel object, indicated by DI > 50. ns p-value > 0.05.

**Supplemental Figure 6:**
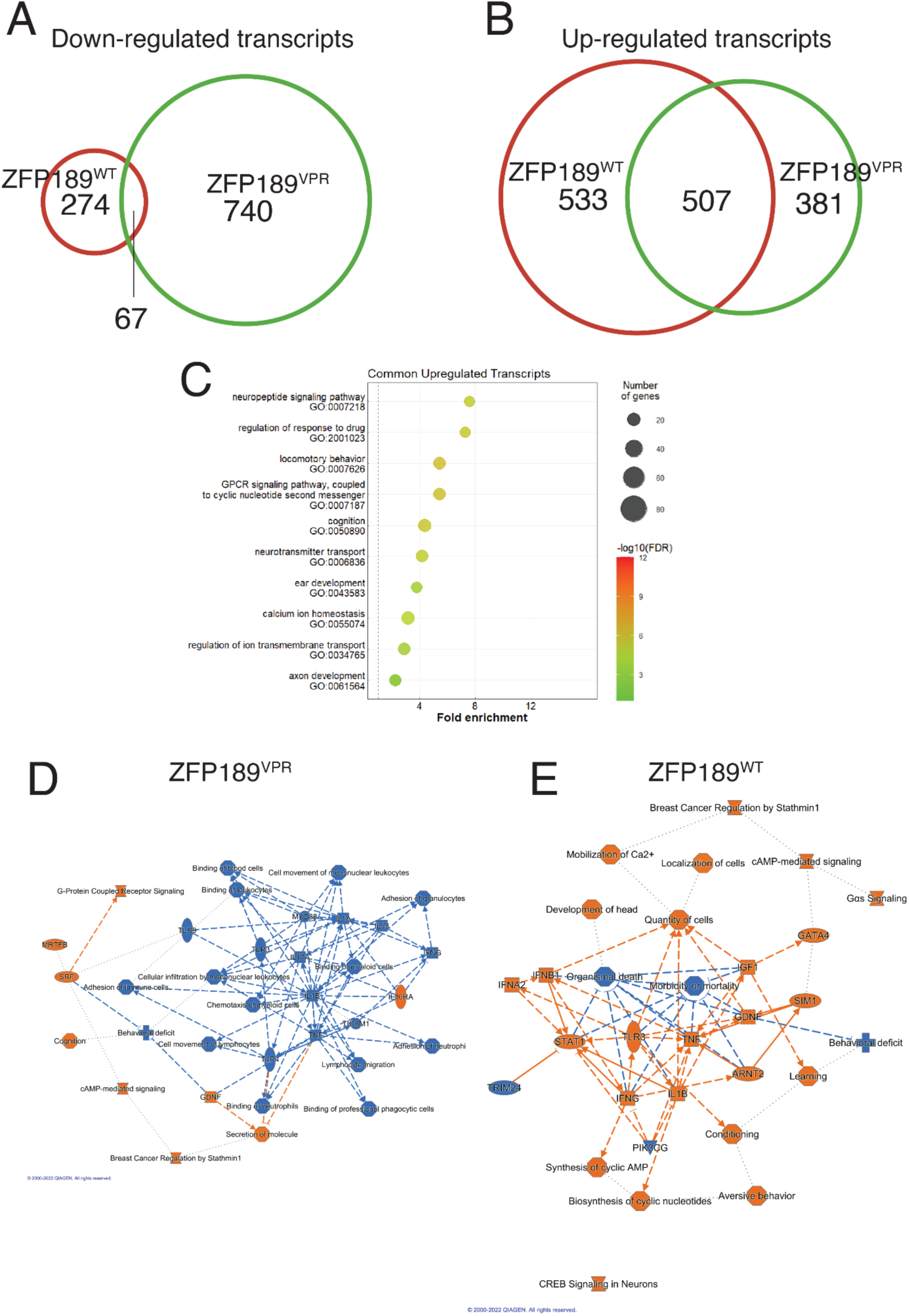
Synthetic ZFP189 TFs have converging up-regulation of PFC transcripts involved in synaptic neurotransmission. Venn diagrams comparing down-**(A)** and up-regulated **(B)** transcripts regulated by each ZFP189 TF. **(C)** Gene-ontology analysis of the 507 commonly up-regulated transcripts from panel B. Ingenuity pathway analysis of ZFP189^VPR^-regulated **(D)** and ZFP189^WT^-regulated (E) transcripts.

**Supplemental Figure 7:**
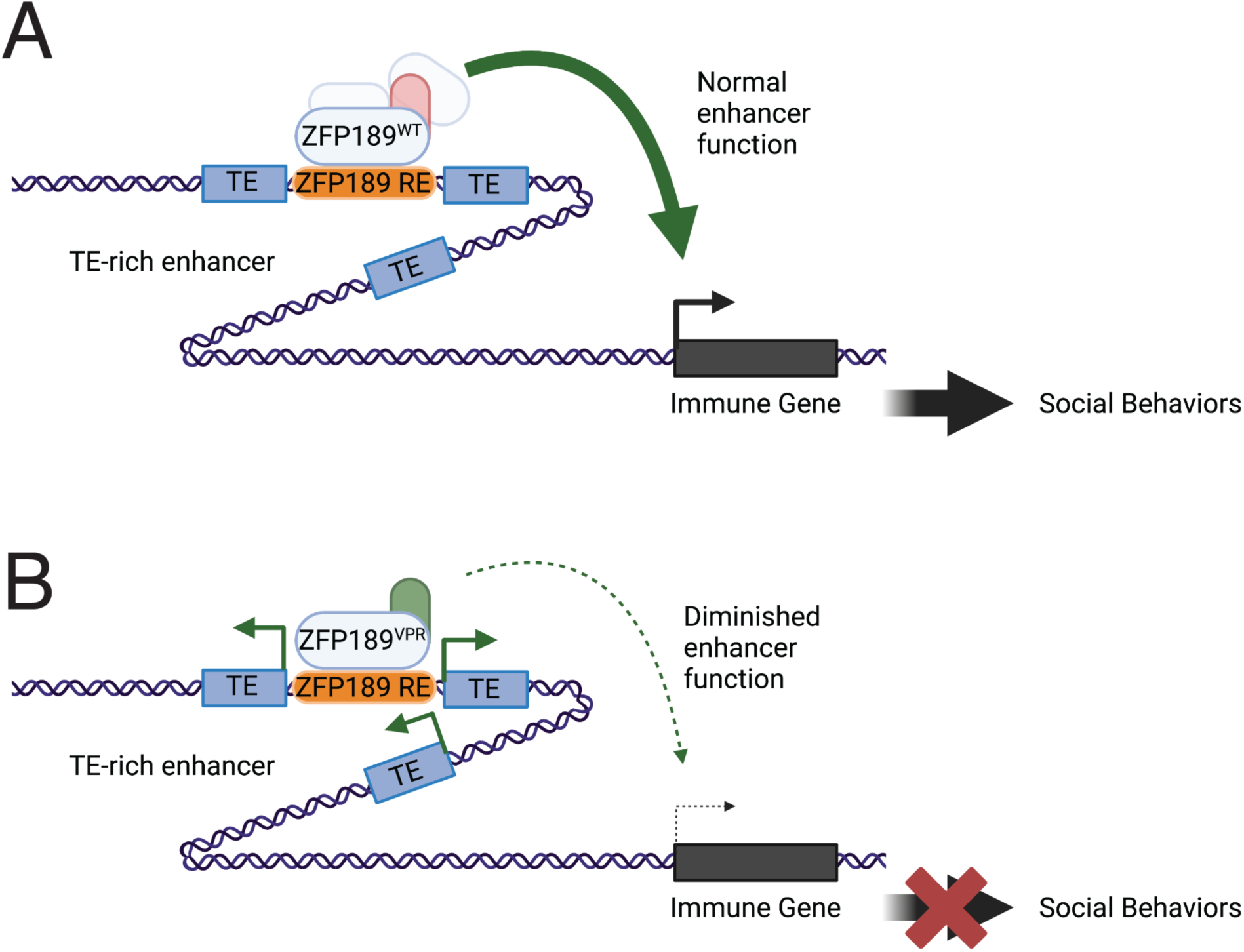
ZFP189-mediated mechanism by which activation of TEs leads to diminished immune transcripts and social behaviors. **(A)** Cartoon representing how the normal function of ZFP189^WT^ can lead to the activation of immune-related transcripts and normal social behaviors. ZFP189^WT^ binds DNA motifs in TE-rich enhancers, represses the transcription of proximal TEs, recruits co-factors, and enables *cis*-regulatory action of this enhancer to activate the expression of immune genes in PFC, which, in turn, promotes social behaviors. **(B)** Alternatively, ZFP189^VPR^ binds ZFP189 DNA motifs in TE-rich enhancers and activates the transcription of proximal TEs, and renders the enhancer nonfunctional. This results in lowered expression of *cis-*regulated immune genes and diminished social behaviors. Cartoon made in Biorender.com.

